# Joint modeling of multi-timepoint spatial observations for time-resolved spatial-unit-specific gene regulatory network inference

**DOI:** 10.64898/2026.07.28.738702

**Authors:** Yibing Jiang, Yurui Li, Qiqi Xie, Yang Li, Haohan Wang

## Abstract

**Background:** How gene regulatory programs reorganize across space and time is central to development and disease, but current methods infer regulatory structure from single snapshots. The emergence of spatiotemporal transcriptomics calls for methods that resolve regulation along both axes.

**Results:** We introduce SpaTemGRN, which jointly models observations from multi-timepoint slides in spatiotemporal transcriptomics to infer time-resolved spatial-unit-specific putative GRNs. Compared with existing methods, SpaTemGRN allows later-stage units to borrow statistical strength from spatially proximate and temporally preceding units. In the *App^NL−G−F^* mouse model of Alzheimer’s disease, SpaTemGRN revealed region- and age-dependent strengthening of complement–glia regulatory coupling. An early, broadly distributed complement signature precedes later, spatially focal coupling with astrocytic and microglial responses. In the developing mouse embryonic brain, SpaTemGRN identified progressively sharpening and spatially segregated regulatory programs as the early neural tube regionalizes. Across simulated datasets, SpaTemGRN recovered regulatory edges more robustly than four existing methods and maintained the most stable performance across stages.

**Conclusions:** SpaTemGRN provides a flexible hypothesis-generating framework for investigating how spatially localized gene–gene dependencies are remodeled across biological stages. All source code for SpaTemGRN is available at https://github.com/yibingjiang/SpaTemGRN.

## 1 BACKGROUND

Gene regulatory programs are not fixed properties of a cell type, but are continually reshaped by tissue position and developmental or pathological stage [1–4]. Resolving this reorganization across space and time is central to understanding organ development and disease progression. Spatiotemporal transcriptomics is beginning to make these dynamics observable by profiling tissue sections across development [5–7], tissue regeneration [8], or disease progression [9–11]. These datasets make it possible to study how gene regulatory networks (GRNs) evolve jointly across both space and time. Realizing that opportunity, however, requires inference methods that can recover regulatory structure as it changes along both axes.

Current GRN inference methods, however, are not designed to jointly exploit this spatiotemporal structure. Traditional GRN methods like GRNBoost2 infer a single global or population-level network from pooled cells [12–14]. With the emergence of single-cell transcriptomic data, many methods have inferred cell-type-level networks by aggregating expression across cells, which obscures cell-level regulatory heterogeneity [15–18]; in a complementary direction, some methods have focused on inferring cell-specific GRNs [19, 20]. In spatial transcriptomics settings, SVGRN, previously proposed by our team, and CeSpGRN further advanced this line by enabling cell-specific GRN inference that accounts for spatial context [21, 22]. Across all of these, however, regulatory variation is captured within a single spatial or developmental snapshot, and temporal progression across stages is not modeled. Consequently, when these methods are applied to spatiotemporal data, each time point must be analyzed separately: a network is inferred independently for every slide and the ordered progression linking successive stages is discarded. Yet the stages profiled in a spatiotemporal experiment are not independent biological states. In both development and progressive disease, the state of a tissue at one stage arises from the state that preceded it: progenitors give rise to their descendants through ordered fate decisions, and pathological burden accumulates on tissue that has already been altered at earlier stages [23–25]. Regulatory programs are therefore remodeled incrementally rather than rebuilt at each stage, so networks at adjacent stages share substantial structure [26]. This continuity is what makes joint modeling informative: an observation at a later stage is more precisely estimated when it can draw on earlier observations along the same progression, whereas per-slide inference must estimate each network from one section alone and discard the shared structure that the progression itself implies.

Therefore, here we introduce SpaTemGRN, a framework that jointly models observations from multi-timepoint slides in spatiotemporal transcriptomics to infer putative spatial-unit-specific GRNs at each time point. In contrast to other spatial-unit-specific methods that must be run separately on each slide, SpaTemGRN estimates shared dependency structure across all time points within a single framework and then specializes it, so that later-stage units borrow statistical strength from spatially proximate and temporally preceding units rather than being estimated in isolation. In a spot-based longitudinal mouse Alzheimer’s disease dataset, SpaTemGRN identified anatomically localized and stage-dependent complement–glia network remodeling. Further, in a bead-level mouse embryonic development dataset, SpaTemGRN recovered spatially organized regulatory programs associated with early brain regionalization. Finally on simulated data, we benchmarked SpaTemGRN against four GRN inference methods [14, 19, 20, 22]. SpaTemGRN achieved the most robust overall edge recovery, with the greatest advantages under higher noise, larger gene sets, and greater cell numbers. It also provided the most stable recovery across temporal stages. SpaTemGRN therefore offers a hypothesis-generating framework for resolving spatiotemporal remodeling of putative regulatory networks from spatiotemporal transcriptomic data.

## 2 RESULTS

### 2.1 Model overview

SpaTemGRN extends our previous spatially varying GRN model to infer time-resolved spatial-unit-specific putative GRNs from spatiotemporal transcriptomics [21]. The model takes as input a gene expression matrix together with each observation’s spatial coordinates and temporal stage (Fig. 1). A structural-equation-model-inspired GRN component is embedded within a conditional variational autoencoder (CVAE): an encoder maps expression to a latent representation conditioned on the joint spatiotemporal context, and a decoder reconstructs expression from that latent variable and context, with a GRN layer and its inverse propagating oriented gene–gene dependencies in both directions. Training proceeds in two stages: stage 1 learns a single global network that captures system-wide dependency structure across all observations and stages; stage 2 then specializes this network to each target observation, refining a spatial-unit-specific GRN using information borrowed from spatially and temporally proximate observations. The resulting spatial-unit-specific edge weights can be compared across tissue space and biological time. The full model formulation, training objectives, and spatiotemporal kernel are described in the Methods section.

**Figure 1.**
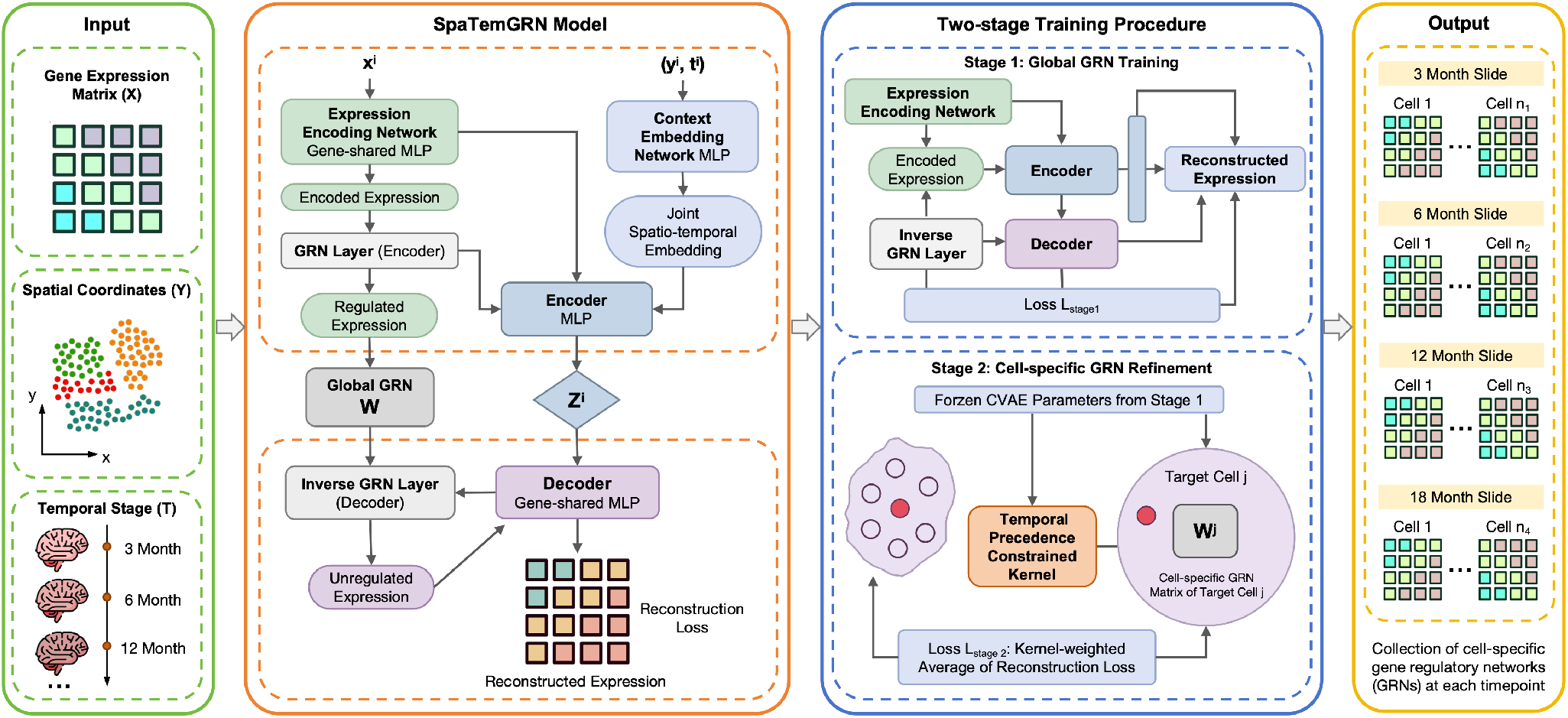
An overview of the SpaTemGRN model architecture, here taking cell-level case as an example. The model takes cell-level gene expression with corresponding spatial coordinates and temporal stage as input. A CVAE encoder maps each cell to a latent representation using both expression and spatiotemporal context, and a decoder reconstructs gene expression conditioned on the same context. A GRN layer (network-structured transformation) and an inverse GRN layer parameterize oriented gene–gene dependencies.

### 2.2 Spatiotemporal dynamics of complement–glia networks in Alzheimer’s disease

In Alzheimer’s disease (AD), a central question is how regulatory programs reorganize across brain regions and disease stages. Amyloid-*β* (A*β*) plaques, tau pathology, and neuroinflammatory responses accumulate in region- and stage-dependent patterns [27, 28]. Chronic microglial activation contributes to disease progression [29, 30], while disease-associated microglia (DAM) cluster around plaques and up-regulate *Trem2, Apoe*, and antigen-presentation genes [31]. Longitudinal and spatial transcriptomic studies document shifts in microglia and astrocytes with plaque burden [25, 32], but do not resolve how gene–gene regulatory dependencies are rewired across space and stage [33]. We therefore used SpaTemGRN to trace when and where AD-associated putative networks rewire in an age-ordered amyloid model.

We analyzed a publicly available spatial transcriptomics dataset of *App*^*NL−G−F*^ knock-in and wild-type mice at 3, 6, 12, and 18 months [11]. The *App*^*NL−G−F*^ model provides an age-ordered system of amyloid-plaque progression and associated glial responses, rather than a tau-based Braak-staging model [34]. The dataset contains 10,327 transcriptomic profiles from 20 coronal brain sections, with approximately 500 profiles per section. Each profile represents a 100-*µ*m-diameter tissue domain integrated with pathological annotations [11]. During preprocessing, we observed an anomalous global reduction in gene expression at 6 months in both genotypes. We therefore applied a spatially guided correction, with the procedure and external validation detailed in Additional file 1. We then selected one representative brain slide per genotype and age, yielding eight slides for analysis^*^. SpaTemGRN inference was restricted to the 56 plaque-induced genes (PIGs) defined in the source study [11]. These genes are consistently elevated in plaque-proximal tissue domains relative to plaque-distal regions and capture plaque-associated inflammatory and glial-response programs [11]. This gene set enabled us to examine spatial and temporal variation in the inferred network structure of plaque-associated immune and glial-response transcripts.

#### Early complement-associated edge patterns suggest amyloid-linked glial priming

We first considered the inferred *C1qa-C4b* edge as a transcript-level dependency between two complement-related genes within the plaque-induced gene panel. In AD, *C1qa-C4b* edge weights were already elevated at 3M and rose modestly from 6–12M in hippocampal and adjacent cortical spots (Fig. 2A and Fig. 2C^†^). This pattern indicates that the spatial-map elevation reflects a broad distributional separation already present at 3M across all three Level 1 brain regions, including the low-pathology brain stem. Meanwhile, Fig. 2D-E shows that the genotype × age interaction effect size was small for nearly all Level 2 brain regions. This suggests that the AD–WT difference in this edge was largely established early and remained relatively stable in magnitude across the age-dependent amyloid-associated trajectory, rather than progressively widening with age. Thus, the temporal increase observed in the spatial maps may reflect geographic expansion of an already-engaged complement-associated transcriptional program rather than a progressively deepening AD-specific divergence. These results suggest that *App*^*NL−G−F*^ tissue contains an early complement-associated transcriptomic network signature that is broadly distributed across brain regions and may reflect an amyloid-associated glial priming state.

**Figure 2.**
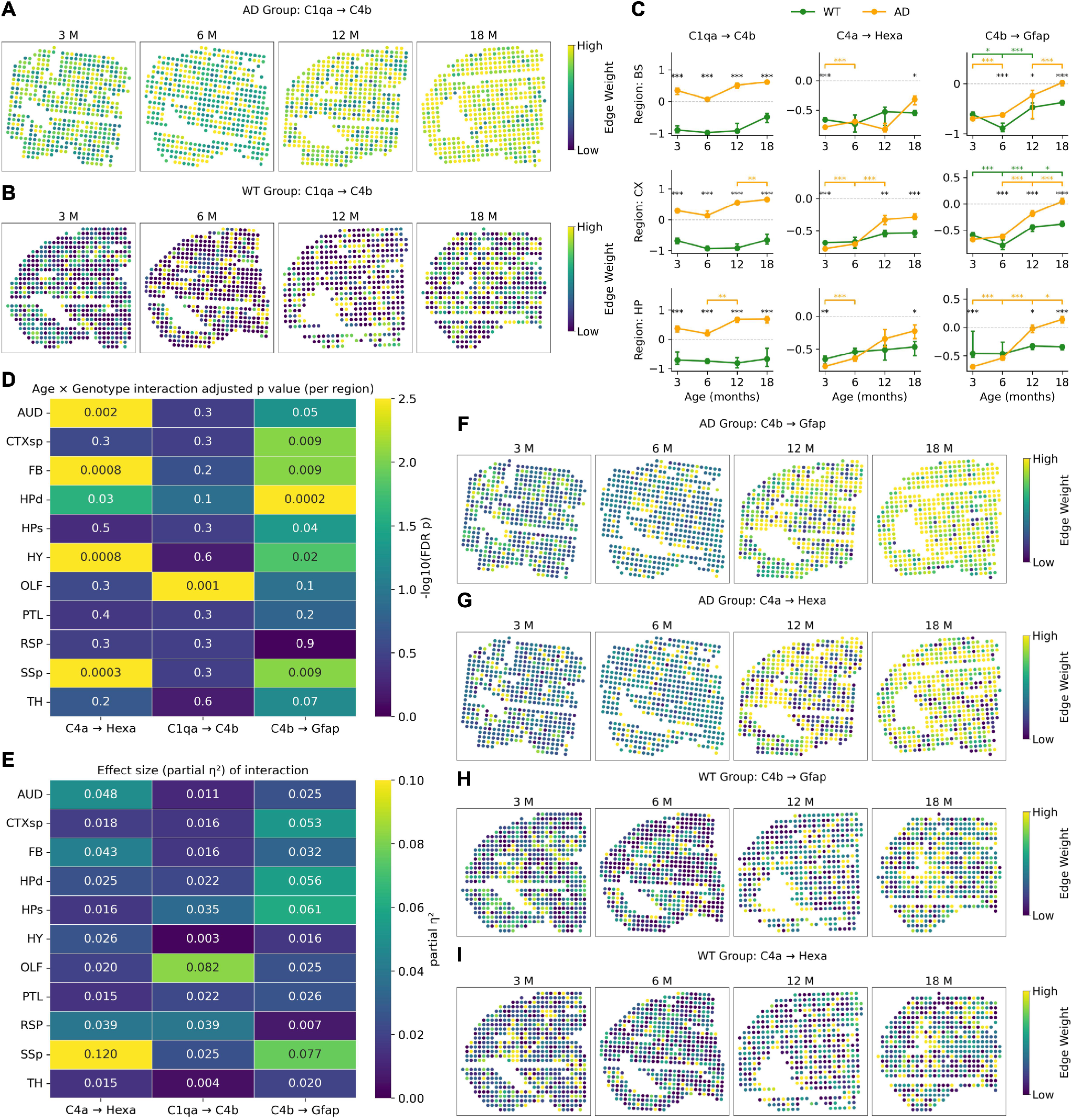
**(A-B)** Spatial edge-weight maps for the inferred *C1qa-C4b* interaction in the (A) AD and (B) WT groups. **(C)** AD-vs-WT comparison of the three complement--glia edges across age and region. Normalized per-spot edge weights for *C1qa-C4b, C4a-Hexa*, and *C4b-Gfap* (columns) within Level 1 regions BS, CX, and HP (rows) at 3, 6, 12, and 18 M, by genotype (WT, green; AD, orange). Points are medians; error bars are 95% bootstrap confidence intervals of the median (1000 resamples). Brackets denote AD-vs-WT contrasts at matched age (black) and within-genotype age contrasts (AD, red; WT, blue), tested by Mann--Whitney *U* with BH-FDR (^*\**^*<*0.05, ^*\*\**^*<*0.01, ^*\*\*\**^*<*0.001). **(D-E)** Per-region genotype *×* age interaction. For each edge (columns) and Level 2 region (rows), heatmaps show the interaction term from a per-region two-way ANOVA on normalized edge weights: (D) *−* log_10_ of the BH-FDR-adjusted *p*-value (*p*_adj_); (E) partial *η*^2^ effect size. Regions COM and ENTI were excluded (insufficient spots for stable estimation). **(F-I)** Spatial maps for the inferred C4-linked glial edges: (F) AD *C4b-Gfap*; (G) AD *C4a-Hexa*; (H) WT *C4b-Gfap*; (I) WT *C4a-Hexa*. In all spatial maps (A-B, F-I), each dot is a spatial transcriptomic spot plotted at its spatial coordinates (x, y), and columns show 3, 6, 12, and 18 months (M).

#### C4-linked glial-response edges strengthen with amyloid-associated progression

We next considered two inferred transcript-level edges linking complement-related genes with glial-response markers: *C4b-Gfap* and *C4a-Hexa*. These edges connect complement-associated transcripts with markers of astrocytic reactivity and microglial lysosomal activity, respectively, and therefore show how complement-related transcriptional programs become co-ordinated with glial responses during disease progression. Spatially, both edges exhibited delayed strengthening relative to the early *C1qa-C4b* edge: AD weights were low at 3M, with distinct hippocampal/subicular and cortical hotspots emerging by 6M and 12M and becoming broadly elevated across hippocampus and large cortical territories by 18M (Fig. 2F-G, Fig. 2C); WT remained low across ages with only minor late-age drift (Fig. 2H-I). Both edges also showed genotype × age interaction effect sizes larger than those for the early *C1qa-C4b* edge (Fig. 2D-E), suggesting that these complement–glial transcript-level dependencies become more AD-associated with age rather than being fully established at the earliest stage.

### 2.3 Spatially structured network patterns in mouse embryonic brain development

Spatial position and biological stage also structure normal brain development. As the embryonic brain forms, spatially restricted gene regulatory programs progressively partition the neural tube into distinct anatomical territories [24]. We therefore used SpaTemGRN to study how regulatory programs are spatially organized across the embryonic brain and how they are remodeled as development proceeds. We used Slide-seq v2 data from Sampath Kumar et al. [6], which provides whole-embryo spatiotemporal transcriptomic maps at near-cellular resolution (∼ 10 *µm* bead diameter) during the onset of organogenesis. We selected one sagittal section per stage from two developmental time points, embryonic day E8.5 and E9.5, during which the neural tube regionalizes into the prosencephalon (forebrain), mesencephalon (midbrain), and rhombencephalon (hindbrain) [6, 35, 36]. After restricting to beads annotated as brain and filtering low-quality beads with low gene detection, we retained 1,032 beads at E8.5 and 2,407 beads at E9.5. Across the 3,439 retained beads, we inferred bead-specific GRNs over a panel of 320 genes selected through a hybrid strategy: differential expression between E8.5 and E9.5, curation from established neural developmental signaling pathways, and spatial variability assessed by Moran’s I within each stage.

#### Bead-specific network profiles form spatially coherent domains

We first studied whether the inferred bead-specific GRNs encode spatially organized regulatory identities. After standardization and PCA, we performed KMeans clustering independently within each developmental stage (*k* = 7). When projected back onto the original tissue coordinates, the GRN-derived clusters formed contiguous spatial domains rather than scattered or intermixed patterns (Fig. 3B). We further compared these GRN-derived clusters with clusters obtained directly from the gene expression matrix using the same number of clusters. Expression-based clustering also recovered spatial structure, particularly at E9.5, but showed more local mixing across neighboring domains (Fig. 3A). In contrast, GRN-based clustering produced broader and more coherent spatial partitions, especially at E9.5, where clusters more clearly separated major anatomical territories along the anterior–posterior axis. This comparison suggests that the inferred bead-specific network profiles preserve spatial organization while capturing edge-weight structure that is not identical to expression-level clustering.

**Figure 3.**
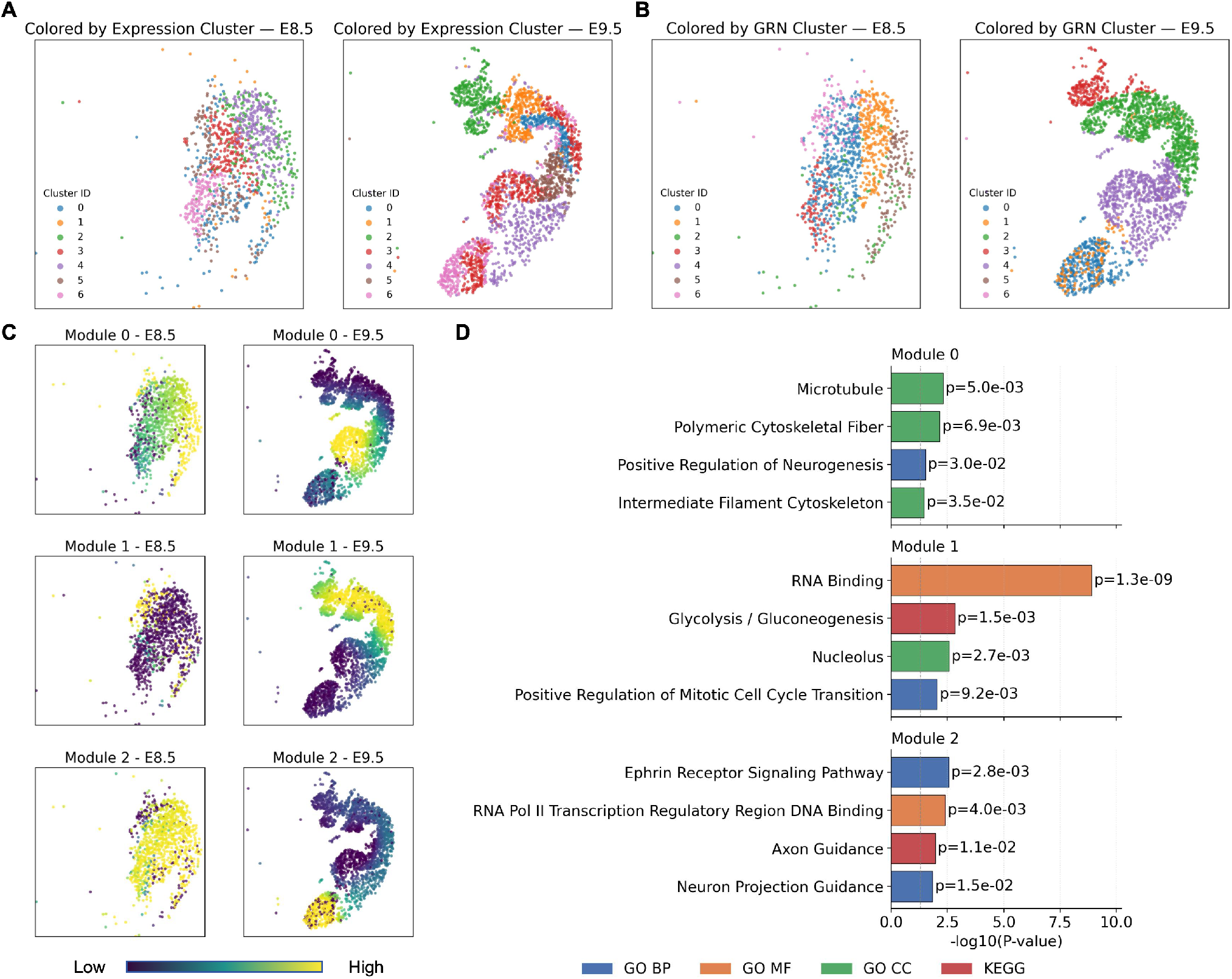
A: Spatial maps of annotated brain-region labels for the selected E8.5 and E9.5 Slide-seq v2 embryo sections. Each point represents one bead and is colored by anatomical annotation. B: Spatial maps of GRN-derived bead clusters at E8.5 and E9.5. Beads were clustered using absolute off-diagonal entries from bead-specific GRN adjacency matrices. C: Spatial activity maps of three edge-based GRN modules at E8.5 and E9.5; module activity was calculated as the mean absolute edge weight of all regulatory edges assigned to each module. D: Enriched GO and KEGG terms for genes participating in each edge-based GRN module; bars show −log10(P-value)

#### Edge-based network modules reveal distinct neural developmental programs

We next explored whether groups of regulatory edges co-vary across space and developmental stage. We constructed an edge-by-bead matrix from the absolute edge weights of all inferred bead-specific GRNs, retained the top 5,000 most spatially variable edges, and clustered them into three modules. For each module, we computed a per-bead activity score as the mean absolute weight of its assigned edges and mapped this score across spatial coordinates at E8.5 and E9.5 (Fig. 3C). The three modules occupied distinct, largely non-overlapping spatial domains, indicating that SpaTemGRN decomposes developmental regulatory structure into spatially localized regulatory programs rather than a single global network. This separation was clearest at E9.5, where each module was active in a different territory of the embryo and the three together tiled complementary domains. At E8.5, the same modules were already spatially structured, but their boundaries were more diffuse and their territories overlapped more extensively than at E9.5. To characterize the biological programs underlying these spatial domains, we performed gene set over-representation analysis on the genes participating in each module’s edges, testing against the GO Biological Process, Molecular Function, and Cellular Component libraries and the KEGG 2019 Mouse library (top enriched terms are shown in Fig. 3D). The three modules corresponded to three distinct regulatory programs. Module 0 was enriched for neuronal cytoskeletal assembly and neurogenesis, driven by neuron-specific tubulins and microtubule-associated proteins (*Tubb3, Map1b*), the neurofilament/intermediate-filament network (*Nefl, Nefm, Ina, Vim*), and the immature-neuron transcription factor *Sox11*. Module 1 was dominated by a biosynthetic and proliferative signature, driven by glycolytic enzymes (*Ldha, Pkm, Eno1, Aldoa*), ribogenesis factors (*Npm1, Ncl, Nop56/58*), cyclin-D cell-cycle regulators (*Ccnd1, Ccnd2*), and anterior neural-progenitor transcription factors (*Sox2, Otx2, Zic1*). Module 2 coupled axon guidance and boundary signaling to regional transcriptional patterning, driven by Eph receptors and ephrins (*Efnb1/2, Epha4/7, Ephb2*) alongside a broad set of positional transcription factors spanning the neural axis (forebrain *Six3*/*Lhx2*, mid-hindbrain *En1*/*Gbx2*, and hindbrain *Hoxb2/3/4*). The three modules describe complementary phases of early neural development: progenitor proliferation (Module 1), post-mitotic neuronal differentiation and neurite assembly (Module 0), and axon guidance coupled to regional patterning (Module 2). Their increasing spatial segregation from E8.5 to E9.5 indicates that SpaTemGRN resolves these programs into progressively more distinct spatial territories as the embryonic brain matures.

### 2.4 Benchmark on simulated datasets with ground-truth networks

We evaluated SpaTemGRN on simulated data with high transcriptional noise and few recovered cells and genes per section in order to mimic real spatial transcriptomic data and determine whether the inferred networks can support biological conclusions. We used simulated data generated by scMultiSim [37], which produces scRNA-seq expression profiles from user-specified, cell-resolved GRNs while preserving the spatial organization of cells. To simulate the temporal axis of spatiotemporal transcriptomics, we extended the simulator with a chain-based design in which spatial-unit-specific GRNs evolve progressively across stages, so that adjacent stages differ incrementally along a shared progression (the full construction is provided in Additional file 1). Within this framework we systematically varied the factors that most affect GRN recovery in practice: the number of genes, the number of cells, and the intrinsic transcriptional noise level, thereby providing a stricter test of robustness [37]. To benchmark GRN inference performance, we adopted the early precision ratio (EPR) evaluation metric from the BEELINE framework [38], which assesses how well an inferred network recovers the ground-truth regulatory edges. Early precision is defined as the proportion of true regulatory edges among the top *k* predicted edges, where *k* is set to the number of edges in the true GRN. EPR normalizes this quantity by the expected early precision under random edge predictions, yielding a scale-free measure of enrichment for true positives in the highest-confidence edge set.

#### Spatial-unit-specific refinement recovers spatial and temporal network variation

Fig. 4A shows a representative simulated layout (three stages, 500 cells, 110 genes, and 0.8 intrinsic noise). Our construction produces ground-truth networks with genuine structure along both axes SpaTemGRN is designed to recover. Within each stage, scMultiSim generates cell-specific GRNs by progressively rewiring regulatory edges from cell to cell, and cells sharing a GRN class are placed in contiguous spatial domains, so that spatially neighboring cells tend to share regulatory structure. Across stages, the networks are chained so that each stage evolves incrementally from its predecessor, mimicking the continuous, ordered progression of a disease axis while preserving spatiotemporal smoothness; temporal precedence holds by construction, since each stage (*t*) is generated using only information from earlier stages (≤ *t*− 1). The detailed simulation procedure (the spatial layering scheme and the stage-to-stage rewiring) is given in Additional file 1.

**Figure 4.**
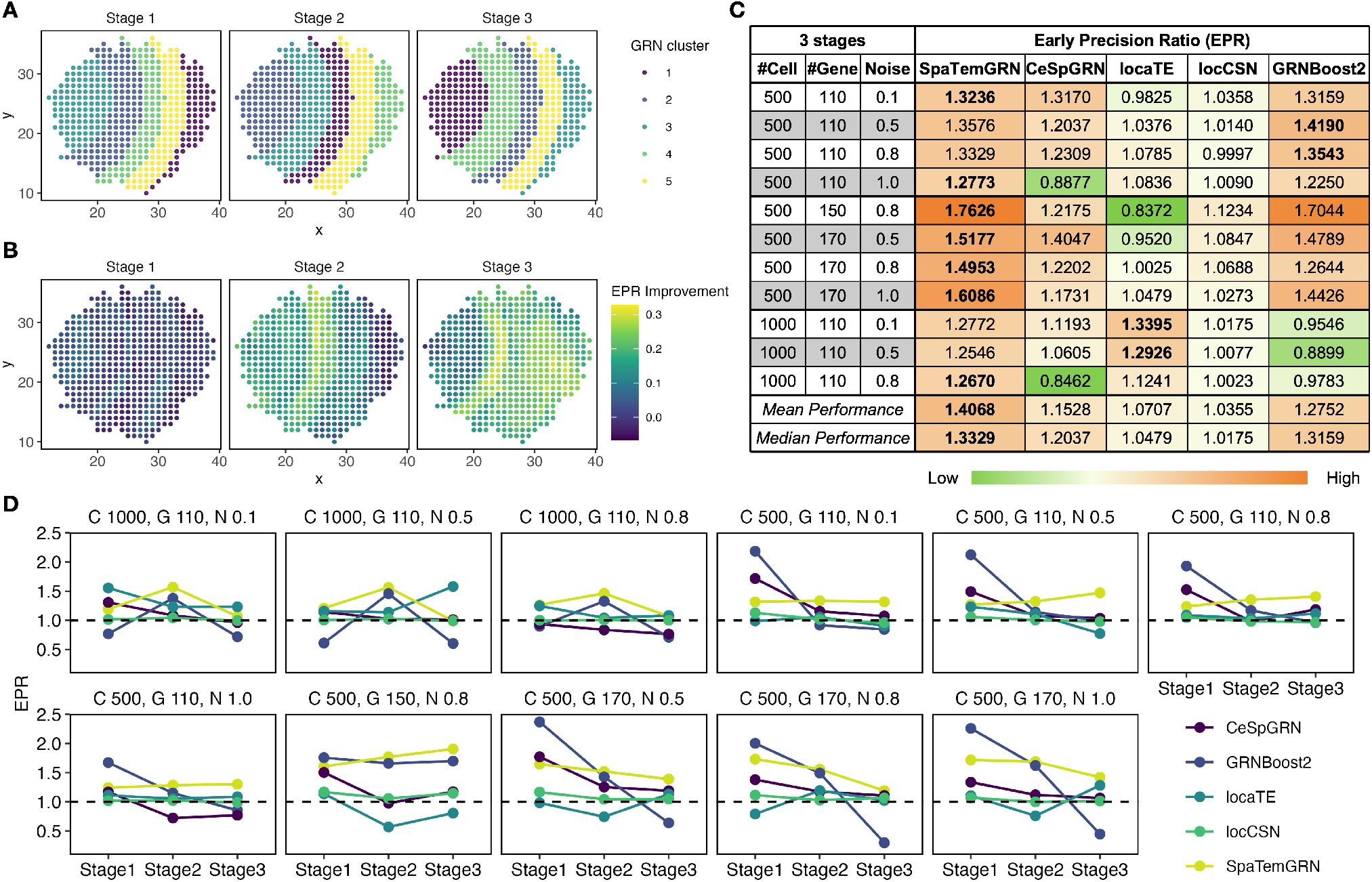
A: The GRNs in all cells of three stages are clustered into five clusters and the cluster ID for each cell is shown in their spatial layout. B: For each cell, the difference between the initial EPR and that after the spatial-unit-specific training is calculated and shown as a heatmap. C: EPR for SpaTemGRN, CeSpGRN, GRNBoost2, locCSN, and locaTE across 11 simulation settings varying cell number (500, 1000), gene number (110, 150, 170), and intrinsic noise level (0.1--1.0). Color intensity reflects metric value from low (dark green) to high (dark orange). D: Stage-wise mean EPR across the 11 simulation settings for all five methods. Each panel corresponds to one setting, defined by the number of cells per stage (C), number of genes (G), and noise level (N).

We then quantified edge recovery for each cell by computing EPR before and after the Stage 2 spatial-unit-specific refinement, starting from the global (whole-tissue) GRN learned in Stage 1. The spatiotemporal distribution of EPR improvement (Fig. 4B) shows broadly positive gains after refinement, indicating that the model fine-tunes GRNs at single-cell resolution rather than producing a single static network. These gains are structured in both space and time. Spatially, EPR gains form banded regions that align with the ground-truth GRN cluster layout: cells belonging to the same simulated GRN cluster show similar improvement, consistent with the model sharing information among spatially proximate cells that have similar regulatory structure. Temporally, EPR improvement increases from Stage 1 to Stage 3, consistent with the role of temporal modeling: later-stage cells borrow information from same-stage spatial neighbors and from earlier-stage cells under the temporal-precedence constraint, stabilizing estimation as regulatory states evolve across stages. SpaTemGRN’s spatial-unit-specific refinement therefore recovers simulated dynamic network rewiring along both spatial and temporal axes.

#### SpaTemGRN shows advantages in more complex settings

We compared our model’s performance to four published GRN inference methods: GRNBoost2, a widely used tree-based regression framework that infers population-level GRNs from gene expression data [14]; locCSN, which constructs cell- and cluster-specific coexpression networks using pairwise independence tests [19]; locaTE, an information-theoretic method that infers cell-specific directed networks from scRNA-seq data [20]; and CeSpGRN, a copula Gaussian graphical method inferring cell-specific GRNs from spatial transcriptomic data [22]. To approximate real-world spatiotemporal transcriptomics, we simulated data spanning the technical and biological variation these methods must tolerate: single-cell and spatial assays carry substantial noise from incomplete RNA capture, batch effects, and biological variability across cell states, compounded in spatial data by tissue handling, sectioning, and barcoding steps that affect capture efficiency and effective resolution [39–41]. Because real datasets also vary widely in gene detection and cell number, robustness across these factors is essential [38, 42–44]. We therefore evaluated all five methods on simulated datasets spanning noise levels of 0.1, 0.5, 0.8, and 1.0, gene counts of 110, 150, and 170, and cell counts of 500 and 1000 per stage. Across all 11 settings (Fig. 4C), SpaTemGRN achieved the highest mean and median EPR and was the only method never to fall below random performance (EPR ≥ 1.0), giving it the highest floor of any method across all conditions. Baselines were competitive only in the simplest setting (fewest cells and genes, low noise); their accuracy declined as noise, gene count, or cell number increased, with GRNBoost2 and CeSpGRN dropping below random under the most challenging conditions. locaTE was the exception, improving with more cells but still degrading at high noise. SpaTemGRN’s advantage was therefore largest in the high-noise, high-dimensional regime that most closely resembles real spatial transcriptomics data.

#### SpaTemGRN achieves more stable GRN recovery across temporal stages

Disease progression and development unfold as ordered stages, so a useful method must recover regulatory structure consistently along that progression: a network recovered well at one stage but lost at the next cannot describe how the program actually changes. To assess this, we computed the stage-wise average EPR for each developmental stage, alongside the overall average across all three stages (Fig. 4D). Although baselines such as CeSpGRN and GRNBoost2 reached competitive or even higher overall mean EPR in some settings, their stage-wise performance was unstable in nearly all of them: both typically showed a sharp drop after Stage 1 or substantial fluctuations across Stages 2 and 3. A favorable pooled average can therefore be misleading for a spatiotemporal task, since a method may score well overall while failing to recover regulatory structure consistently along the progression axis. In contrast, SpaTemGRN yielded much more stable stage-wise EPR and, in many settings, maintained or even improved its EPR from Stage 1 to Stage 3. This stability is consistent with the role of the temporal component: under the temporal-precedence constraint, later-stage cells borrow information from earlier-stage cells, producing reliable inference throughout the trajectory rather than at a single stage only.

## 3 DISCUSSION

Across both an Alzheimer’s disease model and embryonic brain development, SpaTemGRN recovered regulatory programs that reorganize jointly across tissue space and biological progression, resolving structure that single-snapshot methods cannot capture. In the *App*^*NL−G−F*^ brain, SpaTemGRN recovered biologically coherent, stage-dependent edge patterns. The early elevation of the complement-associated *C1qa-C4b* edge is consistent with prior work showing that *C1q* and other complement components rise early in AD models and can localize to synapses before extensive plaque accumulation [45, 46], as well as single-cell studies describing pre-DAM or early activated microglial states with concomitant complement-gene upregulation in amyloid models [25, 31]. In contrast, the delayed strengthening of the *C4b-Gfap* and *C4a-Hexa* glial-response edges is consistent with the neuropathological sequence in which early microglial inflammatory engagement is followed by plaque-associated astrocytic responses [47], and with reported amyloid trajectories in which hippocampal subregions are among the earliest and most heavily affected sites [48, 49]. Therefore, SpaTemGRN recovers a temporal ordering of an early, broadly distributed complement signature followed by later, spatially focal glial-response coupling.

In the developing embryonic brain, SpaTemGRN recovered regionalized regulatory programs in an unsupervised manner. The stronger spatial consolidation of GRN-derived clusters at E9.5 is consistent with known regionalization of the anterior neural tube during this developmental window, when the embryonic brain becomes organized into prosencephalic, mesencephalic, and rhombencephalic territories with regionally patterned transcriptional programs [6, 35, 36]. The progressive sharpening of edge-based module activity from E8.5 to E9.5 parallels the transition from an earlier, loosely organized neural tube toward more clearly delineated compartments, and supports the idea that SpaTemGRN captures the network remodeling that accompanies progressive regionalization [6, 35]. The functional identities of the three edge modules align with the known cellular composition of the developing neural tube, supporting the biological plausibility of the recovered programs. The cytoskeletal and neurogenesis signature of Module 0, together with the immature-neuron factor *Sox11*, which drives neuronal differentiation and neurite outgrowth [50, 51], is consistent with post-mitotic neurons actively extending neurites. The glycolytic and biosynthetic signature of Module 1 reflects the aerobic glycolysis that characterizes dividing neural progenitors and is downregulated as they differentiate [52], consistent with a proliferating, forebrain-biased progenitor pool marked by *Sox2, Otx2*, and *Zic1*. In Module 2, the Eph receptors and ephrins mediate rhombomere-boundary formation and axon guidance in the hindbrain through bidirectional signaling [53], while the accompanying positional transcription factors (*Six3*/*Lhx2, En1*/*Gbx2, Hoxb2/3/4*) establish anteroposterior regional identity across the developing brain [24, 35].

On simulated data, jointly modeling spatiotemporal structure yielded the most robust edge recovery and the most stable performance across stages. The stability of stage-wise performance is particularly relevant for studying progressive biology. Regulatory networks undergo substantial rewiring as biological processes unfold rather than remaining fixed over time, and AD pathology in particular evolves across stages and brain regions rather than appearing uniformly [25, 45, 48]. In mouse AD models, complement activation and microglia-mediated synaptic pathology emerge early, before overt plaque deposition, and plaque burden and microglial responses then expand and intensify across later ages and brain regions [25, 45, 48]. Therefore, a method that recovers regulatory structure well at the earliest stage but deteriorates later is poorly suited to studying disease progression, regardless of how strong its pooled performance appears, while the spatiotemporal kernel and temporal-precedence component of SpaTemGRN are designed precisely to stabilize inference along such a progression axis.

The dominant operational consideration when applying SpaTemGRN is computational cost, which is driven primarily by the number of spatial units rather than by the size of the gene set. On our simulated benchmarks (three temporal stages), a complete two-stage run (global network training followed by spatial-unit-specific refinement) required approximately 17 hours for 500 cells per stage with 110 genes and 21 hours for 500 cells per stage with 170 genes, but 128 hours for 1,000 cells per stage with 110 genes, using four NVIDIA GH200 GPUs on a single node. Increasing the gene set roughly 1.5-fold raised runtime by only about 24%, whereas doubling the number of cells per stage increased runtime by roughly 7.5-fold, indicating that runtime scales steeply and super-linearly with the total number of spatial units while remaining comparatively insensitive to gene-set size (noise level had no appreciable effect on runtime). In practice this means that gene panels can be expanded at modest cost, but that analyses of sections containing many thousands of spatial units require substantial GPU resources and wall-clock time; users working with large tissue sections may wish to restrict inference to an informative gene panel and, where appropriate, to a representative subregion or a down-sampled set of units.

Our method has two main limitations. First, because SpaTemGRN does not require TF–motif or chromatin evidence as input, the inferred edges represent oriented statistical gene–gene dependencies; they should be treated as hypotheses for further testing rather than established regulatory links. Second, inference is computationally intensive: as detailed above, runtime scales steeply with the number of spatial units, which may limit its immediate applicability to atlas-scale datasets. Future work should strengthen the causal interpretation of inferred edges through integration with perturbational data, chromatin accessibility, TF-motif evidence, and experimental validation, and should improve computational scalability to make inference on atlas-scale spatial transcriptomics practical.

## 4 CONCLUSIONS

SpaTemGRN infers spatial-unit-specific putative gene regulatory networks across tissue space and biological time, resolving regulatory structure at the locations and stages at which programs change during development and disease progression. In *App*^*NL−G−F*^ mouse data, SpaTemGRN identified region- and age-dependent complement– glia network patterns. In embryonic mouse brain data, it resolved increasingly distinct spatial programs during early neural-tube regionalization. SpaTemGRN also showed robust edge recovery and stable stage-wise performance across simulated datasets. These results show that SpaTemGRN can generate testable hypotheses about spatiotemporal variation in oriented gene–gene dependencies.

## 5 METHODS

### 5.1 Modeling context-dependent gene--gene dependencies across space and time

To model regulatory relationships that may differ across tissue location and biological stage, we build on the structural equation modeling (SEM) formulation adopted by SVGRN for inferring spatially varying gene regulatory networks (GRNs) from spatial transcriptomics [21, 54]. Let X ∈ ℝ^*n×m*^ denote the gene expression matrix for *n* cells (or more broadly, spatial observations) and *m* genes, 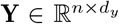 the spatial coordinates (typically *d*_*y*_ = 2), and W ∈ ℝ^*m×m*^ the GRN adjacency matrix, where *W*_*ij*_ parameterizes an inferred oriented gene–gene dependence from gene *i* to gene *j* within the model. Let Z ∈ ℝ^*n×m*^ denote latent variation and noise. In the basic linear SEM, gene expression is represented as the combination of network-mediated gene–gene dependencies and residual latent variation:

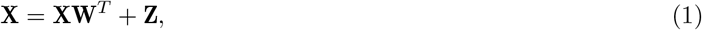

so that Z = X(I −W^*T*^ ) and, when (I −W^*T*^ ) is invertible, X = Z(I −W^*T*^ )^*−*1^. These equivalent forms motivate a GRN “layer” in which (I −W^*T*^ ) and (I −W^*T*^ )^*−*1^ encode how inferred gene–gene dependencies are propagated in the model.

Because regulatory relationships may be nonlinear and may vary across local tissue environments, SVGRN relaxes the linearity assumption in Eq. 1 by replacing the linear mappings with learnable nonlinear functions *f* and conditioning these functions on spatial coordinates Y [21, 55]. Here, conditioning means providing Y as additional inputs so that the learned mappings between latent variation and gene expression can vary with the spatial location of each observation. Concretely, SVGRN writes:

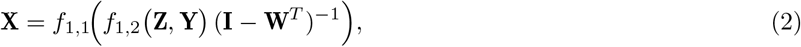

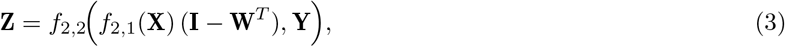

where *f*_1,1_, *f*_1,2_, *f*_2,1_, *f*_2,2_ are neural networks and the (I −W^*T*^ ) / (I −W^*T*^ )^*−*1^ transformations integrate the GRN structure into the generative (noise-to-expression) and inference (expression-to-noise) directions [21, 55]. Constraints and regularization on W used to ensure a well-defined model are handled in the training subsection (see Section 5.3).

The above formulation is designed for single-timepoint spatial transcriptomics, where regulatory variation is modeled across space but not across ordered biological stages. Spatiotemporal transcriptomic datasets, in contrast, profile tissues across developmental, disease, or treatment stages and therefore provide an additional biological axis along which putative regulatory relationships may be remodeled. To exploit this richer data structure, we extend Eqs. 2 and 3 to incorporate temporal stage. Let T ∈ ℝ^*n×*1^ denote a temporal variable for each cell (e.g., experimental time point, disease stage, or pseudo-time); when T is discrete, it is encoded into a suitable numeric representation for conditioning. We condition the same SEM mappings on the joint spatiotemporal context (Y, T) by augmenting the conditioning inputs with T:

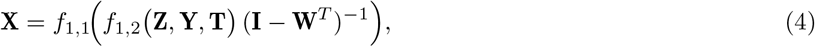

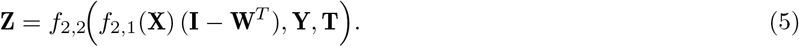

Conditioning on both Y and T allows SpaTemGRN to represent local spatial heterogeneity and stage-dependent remodeling of inferred regulatory structure within a unified SEM. The spatiotemporal training procedure that yields spatial-unit-specific GRNs, including the temporal precedence constraint used when borrowing information across cells, is described in Section 5.3.

### 5.2 A latent-variable framework for spatiotemporal putative regulatory network inference

We implement the spatiotemporal SEM in Eqs. 4 and 5 using a conditional variational autoencoder (CVAE). The CVAE provides a nonlinear latent-variable model for gene expression in which each spatial unit *i* has observed expression x^*i*^ ∈ ℝ^*m*^, spatial coordinates 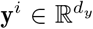, and temporal stage *t*^*i*^ ∈ ℝ. The encoder (recognition network) *q*_*ϕ*_(z^*i*^ | x^*i*^, y^*i*^, *t*^*i*^) maps the observed expression to a variational posterior over latent variation and noise z^*i*^ ∈ ℝ^*m*^, conditioned on (y^*i*^, *t*^*i*^). The decoder *p*_*θ*_(x^*i*^| z^*i*^, y^*i*^, *t*^*i*^) reconstructs gene expression from the latent variable and the same spatiotemporal context. As in SVGRN, we use a standard Gaussian prior *p*(z^*i*^) = *N* (0, I) to encourage a structured and continuous latent space [21].

The model architecture mirrors the SEM transformations by explicitly incorporating the GRN adjacency matrix W into both the expression-to-latent and latent-to-expression mappings. We first embed the spatiotemporal context by concatenating (y^*i*^, *t*^*i*^) and passing it through a multilayer perceptron (MLP) *E*_*Y T*_ to obtain a low-dimensional context embedding. In parallel, the expression vector x^*i*^ is processed by a gene-shared MLP *E*_*X*_ to produce a gene representation 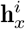. Within the encoder, we apply a network-structured transformation that transforms 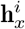 by right-multiplication with (I − W^*T*^ ), yielding a network-propagated representation that is then combined with the context embedding and mapped to the parameters of a Gaussian variational posterior:

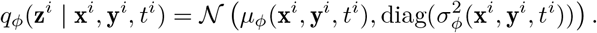

The decoder follows the inverse direction of the SEM: a latent sample z^*i*^ is first combined with the spatiotemporal embedding from *E*_*Y T*_ and passed through a decoding MLP, then transformed by right-multiplication with (I − W^*T*^ )^*−*1^, and finally mapped through a gene-shared output MLP *D* to produce the reconstructed expression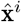.

We train the model by optimizing *ϕ, θ*, and W using the conditional evidence lower bound (ELBO). Let X ∈ ℝ^*n×m*^, 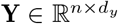, and T ∈ ℝ^*n×*1^ denote the stacked expression, spatial, and temporal variables across all*n* observations, and let Z ∈ ℝ^*n×m*^ denote the corresponding latent variables. The objective is

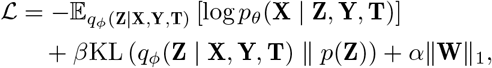

where the first term is the negative conditional reconstruction log-likelihood, the second term is the Kullback– Leibler (KL) divergence regularizing the variational posterior toward the prior, *β* controls the strength of the KL regularization, and *α*∥W∥_1_ promotes sparsity in the inferred GRN.

### 5.3 Learning global and spatial-unit-specific putative regulatory networks

SpaTemGRN learns putative regulatory networks in two stages. Stage 1 learns a global network matrix W that captures system-wide gene–gene dependency structure across all observations and stages. Stage 2 then specializes W to each target observation by refining a spatial-unit-specific network W^*j*^ using information borrowed from spatially and temporally proximate observations, while enforcing a temporal precedence constraint.

In the first stage, we train the CVAE on all observations to obtain an initial system-level GRN (or global GRN). The model is conditioned on each observation’s spatiotemporal context c^*i*^ = (y^*i*^, *t*^*i*^). The Stage 1 objective is the averaged negative ELBO with an *L*_1_ sparsity penalty on W:

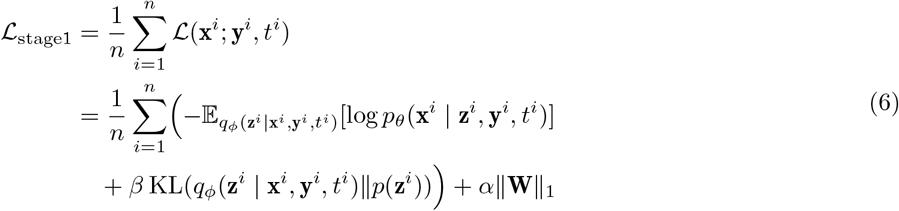

where the first term is the conditional reconstruction loss, the second term regularizes the variational posterior toward the prior (weighted by *β*), and *α* ∥W∥ _1_ promotes sparsity in the global GRN. This stage provides a stable network initialization before estimating spatial-unit-specific regulatory variation. To keep (I − W^*T*^ ) well conditioned during training, we impose two constraints on W: (i) diagonal entries are fixed to zero, excluding self-edges from the inferred network for identifiability; and (ii) an L1 sparsity penalty shrinks off-diagonal weights toward zero, which empirically keeps the spectral radius of W below one and supports stable inversion of (I − W^*T*^ ). We do not impose a hard acyclicity constraint, such as the continuous DAG penalty used in prior work [55]; instead, we use sparsity regularization to encourage sparse and well-conditioned networks.

The second stage specializes the model to infer spatial-unit-specific GRNs based on the global GRN inferred from stage 1. This refinement is based on three biological assumptions linking spatial location, temporal stage, and gene expression. First, we adopt a biological-progression continuity principle: ordered stages in development, disease progression, or treatment response can often be treated as samples along a progression axis [23]. In AD, this assumption is supported in human studies by molecular changes associated with pathological stage, including Braak-stage-associated transcriptional differences [33]. In amyloid mouse models, including the type of model analyzed in this study, the relevant temporal axis is age-dependent amyloid plaque accumulation and the accompanying evolution of glial states rather than tau-based Braak staging [25]. Rather than treating time points as independent biological states, this principle supports integrating tissue sections from ordered stages into a shared progression axis along which regulatory-network dynamics can be analyzed.

Second, we assume that nearby observations in physical space and adjacent biological stages have more similar putative regulatory architectures than distant observations. In spatial transcriptomics, gene expression patterns often vary smoothly across tissue space because of localized morphogen gradients, cell–cell communication, anatomical organization, and diffusion-limited signaling [56, 57]. Similarly, during ordered biological processes such as development or disease progression, molecular states and their regulatory dependencies are often remodeled gradually over time [26]. Cells proximate in the combined spatiotemporal manifold are therefore expected to share similar regulatory networks, allowing the model to borrow statistical strength from biologically relevant neighbors. Third, we impose a temporal precedence constraint: when refining the GRN at a target stage *t*^*j*^, information used for refinement is restricted to observations from the same or earlier stages (*t*^*i*^ ≤ *t*^*j*^), reflecting the directional ordering of biological time [26, 58].

To implement the second and third assumptions, we introduce a spatiotemporal kernel weight into the loss function. This weight determines how much each observation *i* contributes to the refinement of a target observation *j* based on spatial proximity and temporal ordering. Let 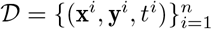our spatiotemporal dataset, where x^*i*^ ∈ ℝ^*m*^ is the gene expression vector, y^*i*^ = (*u*^*i*^, *v*^*i*^) ∈ ℝ^2^ is the spatial-coordinate vector, and *t*^*i*^ ∈ ℝ is the temporal coordinate (age) for spatial unit *i*. We define the temporal-precedence-constrained kernel weight *K*_*ij*_ between observation *i* and target observation *j* as:

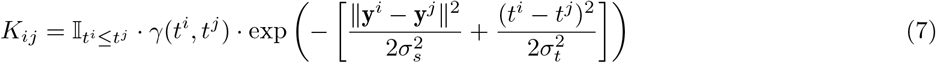

where *σ*_*s*_ and *σ*_*t*_ serve as bandwidth hyperparameters for spatial and temporal decay, respectively, and 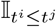 enforces temporal precedence. The factor *γ*(*t*^*i*^, *t*^*j*^) modulates the relative contribution of past-stage versus same-stage cells:

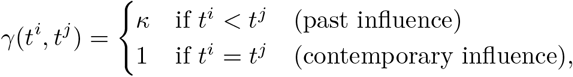

with *κ* ≥ 1 controlling the relative contribution of earlier-stage observations. This reflects the assumption that earlier-stage observations provide information about the trajectory leading to the target stage, whereas same-stage neighbors primarily capture spatial heterogeneity at the current stage. Because same-stage patterns are also partially represented through conditioning on the target context, up-weighting earlier stages helps preserve temporal information in the refinement objective.

For each target observation *j*, we refine W^*j*^ by freezing the shared model parameters (*ϕ, θ*) and optimizing only the target-specific GRN matrix W^*j*^ under a kernel-weighted objective. During this refinement, we condition the model on the target context (y^*j*^, *t*^*j*^), so that W^*j*^ represents the inferred GRN associated with the target observation’s spatial location and temporal stage. The Stage 2 objective is:

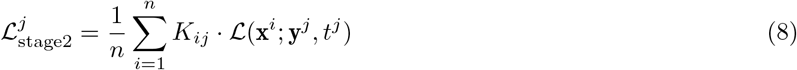

where *K*_*ij*_ is defined in Eq. 7 and ℒ (x^*i*^; y^*j*^, *t*^*j*^) denotes the per-observation negative ELBO (reconstruction plus KL) evaluated for expression x^*i*^ when the model is conditioned on the fixed target context (y^*j*^, *t*^*j*^). By optimizing W^*j*^ to minimize 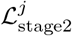, the model borrows information from spatially and temporally proximate observations (via *K*_*ij*_) while respecting the temporal precedence constraint.

Therefore, under the spatiotemporal smoothness assumption, observations proximate to *j* are assumed to have similar expression-associated network structure; consequently, their expression profiles x^*i*^ should be approximately reproducible by a decoder conditioned on the target context (y^*j*^, *t*^*j*^) and governed by W^*j*^ . The overall procedure first learns a global GRN via Stage 1, then iterates over all target observations *j* and refines a spatial-unit-specific GRN W^*j*^ using the kernel-weighted objective in Eq. 8. This yields a collection of spatial-unit-specific GRNs 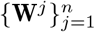 that capture regulatory changes across space and biological time.

## DATA AND CODE AVAILABILITY

The Alzheimer’s disease (AD) spatial transcriptomics dataset used for GRN inference was obtained from the Gene Expression Omnibus (GEO, https://www.ncbi.nlm.nih.gov/geo/) under accession GSE152506. We analyzed four AD slides (N02_D1, N07_C1, N06_D2, N04_E1) and four wild-type (WT) slides (B02_D1, B07_C2, B06_E1, B04_D1). The external AD dataset used as a reference for batch-effect correction was also obtained from GEO under accession GSE174321. The mouse embryo spatiotemporal transcriptomic dataset is available from CEL-LxGENE at https://cellxgene.cziscience.com/collections/d74b6979-efba-47cd-990a-9d80ccf29055; the two sections used in the analysis are 201104_14 (E8.5) and 201104_33 (E9.5). All source code for the SpaTemGRN model and processed data used for analyses are available at https://github.com/yibingjiang/SpaTemGRN.

## ADDITIONAL FILES

### Additional file 1 — supplementary materials

Construction of the simulated spatiotemporal datasets (S1); spatial-guided latent batch correction (S2); and supplementary figures.

## Footnotes

* AD and WT samples were modeled separately and comparisons between genotypes were therefore performed downstream on the inferred edge-weight distributions.

† For (C-E), because one representative section was analyzed per genotype and age, these spot-level tests are intended as exploratory summaries of within-section patterns rather than population-level inference.

## REFERENCES

[1] Nicholas M. Luscombe et al. “Genomic analysis of regulatory network dynamics reveals large topological changes”. In: Nature 431.7006 (2004), pp. 308–312.

[2] Amr Ahmed and Eric P. Xing. “Recovering time-varying networks of dependencies in social and biological studies”. In: Proc. Natl. Acad. Sci. U.S.A. 106.29 (2009), pp. 11878–11883.

[3] Tsukasa Kouno et al. “Temporal dynamics and transcriptional control using single-cell gene expression analysis”. In: Genome Biol. 14.10 (2013), R118.

[4] Cong Ma et al. “Belayer: Modeling discrete and continuous spatial variation in gene expression from spatially resolved transcriptomics”. In: Cell Syst. 13.10 (2022), 786–797.e13.

[5] Yanting Wu et al. “A spatiotemporal transcriptomic atlas of mouse placentation”. In: Cell Discov. 10.1 (2024), p. 110.

[6] Abhishek Sampath Kumar et al. “Spatiotemporal transcriptomic maps of whole mouse embryos at the onset of organogenesis”. In: Nat. Genet. 55.7 (2023), pp. 1176–1185.

[7] Jiexue Pan et al. “Spatiotemporal transcriptome atlas of human embryos after gastrulation”. In: Nature 654.8119 (2026), pp. 751–761.

[8] Jiangshan Xu et al. “A spatiotemporal atlas of mouse liver homeostasis and regeneration”. In: Nat. Genet. 56.5 (2024), pp. 953–969.

[9] Emily Miyoshi et al. “Spatial and single-nucleus transcriptomic analysis of genetic and sporadic forms of Alzheimer’s disease”. In: Nat. Genet. 56.12 (2024), pp. 2704–2717.

[10] Hongyoon Choi et al. “Spatiotemporal characterization of glial cell activation in an Alzheimer’s disease model by spatially resolved transcriptomics”. In: Exp. Mol. Med. 55.12 (2023), pp. 2564–2575.

[11] Wei-Ting Chen et al. “Spatial transcriptomics and in situ sequencing to study Alzheimer’s disease”. In: Cell 182.4 (2020), 976–991.e19.

[12] Sara Aibar et al. “SCENIC: single-cell regulatory network inference and clustering”. In: Nat. Methods 14.11 (2017), pp. 1083–1086.

[13] Carmen Bravo González-Blas et al. “SCENIC+: single-cell multiomic inference of enhancers and gene regulatory networks”. In: Nat. Methods 20.9 (2023), pp. 1355–1367.

[14] Thomas Moerman et al. “GRNBoost2 and Arboreto: efficient and scalable inference of gene regulatory networks”. In: Bioinformatics 35.12 (2019), pp. 2159–2161.

[15] Russell Littman et al. “SCING: Inference of robust, interpretable gene regulatory networks from single cell and spatial transcriptomics”. In: iScience 26.7 (2023), p. 107124.

[16] Lei Hu et al. “STARNet enables spatially resolved inference of gene regulatory networks from spatial multi-omics data”. In: bioRxiv (2025).

[17] Zhanhe Chang et al. “Single-cell and spatial multiomic inference of gene regulatory networks using SCRIPro”. In: Bioinformatics 40.7 (2024), btae466.

[18] Yao Li et al. “SpaGRN: Investigating spatially informed regulatory paths for spatially resolved transcriptomics data”. In: Cell Syst. 16.4 (2025), p. 101243.

[19] Xuran Wang, David Choi, and Kathryn Roeder. “Constructing local cell-specific networks from single-cell data”. In: Proc. Natl. Acad. Sci. U. S. A. 118.51 (2021), e2113178118.

[20] Stephen Y Zhang and Michael P H Stumpf. “Learning cell-specific networks from dynamics and geometry of single cells”. In: Cell Syst. 16.10 (2025), p. 101399.

[21] Yurui Li, Jin Chen, and Haohan Wang. “Spatially varying cell-specific gene regulation network inference”. In: bioRxiv (2025).

[22] Ziqi Zhang et al. “CeSpGRN: inferring cell-specific gene regulatory networks from single-cell multi-omics and spatial data”. In: Bioinformatics 42.6 (2026).

[23] Cole Trapnell et al. “The dynamics and regulators of cell fate decisions are revealed by pseudotemporal ordering of single cells”. In: Nat. Biotechnol. 32.4 (2014), pp. 381–386.

[24] Clemens Kiecker and Andrew Lumsden. “Compartments and their boundaries in vertebrate brain development”. In: Nature Reviews Neuroscience 6.7 (2005), pp. 553–564.

[25] Carlo Sala Frigerio et al. “The major risk factors for Alzheimer’s disease: Age, sex, and genes modulate the microglia response to Aβ plaques”. In: Cell Rep. 27.4 (2019), 1293–1306.e6.

[26] Mukesh Bansal et al. “How to infer gene networks from expression profiles”. In: Mol. Syst. Biol. 3.1 (2007), p. 78.

[27] H Braak and E Braak. “Neuropathological stageing of Alzheimer-related changes”. In: Acta Neuropathol. 82.4 (1991), pp. 239–259.

[28] H Braak and E Braak. “Staging of Alzheimer’s disease-related neurofibrillary changes”. In: Neurobiol. Aging 16.3 (1995), 271–8, discussion 278–84.

[29] Chao Gao et al. “Microglia in neurodegenerative diseases: mechanism and potential therapeutic targets”. In: Signal Transduct. Target. Ther. 8.1 (2023), p. 359.

[30] Aniket Kakkar et al. “Neuroinflammation and Alzheimer’s disease: Unravelling the molecular mechanisms”. In: J. Alzheimers. Dis. 108.1 (2025), pp. 19–41.

[31] Hadas Keren-Shaul et al. “A unique microglia type associated with restricting development of Alzheimer’s disease”. In: Cell 169.7 (2017), 1276–1290.e17.

[32] Shengran Wang et al. “Gene interactions analysis of brain spatial transcriptome for Alzheimer’s disease”. In: Genes Dis. 11.6 (2024), p. 101337.

[33] Jorge L Del-Aguila et al. “A single-nuclei RNA sequencing study of Mendelian and sporadic AD in the human brain”. In: Alzheimers. Res. Ther. 11.1 (2019), p. 71.

[34] Takashi Saito et al. “Single App knock-in mouse models of Alzheimer’s disease”. In: Nat. Neurosci. 17.5 (2014), pp. 661–663.

[35] W Wurst and L Bally-Cuif. “Neural plate patterning: upstream and downstream of the isthmic organizer”. In: Nat. Rev. Neurosci. 2.2 (2001), pp. 99–108.

[36] C D Stern. “Initial patterning of the central nervous system: how many organizers?” In: Nat. Rev. Neurosci. 2.2 (2001), pp. 92–98.

[37] Hechen Li et al. “scMultiSim: simulation of single-cell multi-omics and spatial data guided by gene regulatory networks and cell-cell interactions”. In: Nat. Methods 22.5 (2025), pp. 982–993.

[38] Aditya Pratapa et al. “Benchmarking algorithms for gene regulatory network inference from single-cell transcriptomic data”. In: Nat. Methods 17.2 (2020), pp. 147–154.

[39] Philipp Angerer et al. “Single cells make big data: New challenges and opportunities in transcriptomics”. In: Curr. Opin. Syst. Biol. 4 (2017), pp. 85–91.

[40] Monika Piwecka, Nikolaus Rajewsky, and Agnieszka Rybak-Wolf. “Single-cell and spatial transcriptomics: deciphering brain complexity in health and disease”. In: Nat. Rev. Neurol. 19.6 (2023), pp. 346–362.

[41] Lambda Moses and Lior Pachter. “Museum of spatial transcriptomics”. In: Nat. Methods 19.5 (2022), pp. 534–546.

[42] Yue You et al. “Systematic comparison of sequencing-based spatial transcriptomic methods”. In: Nat. Methods 21.9 (2024), pp. 1743–1754.

[43] Jiao Cao et al. “Decoder-seq enhances mRNA capture efficiency in spatial RNA sequencing”. In: Nat. Biotechnol. 42.11 (2024), pp. 1735–1746.

[44] Jong Kyoung Kim et al. “Characterizing noise structure in single-cell RNA-seq distinguishes genuine from technical stochastic allelic expression”. In: Nat. Commun. 6.1 (2015), p. 8687.

[45] Soyon Hong et al. “Complement and microglia mediate early synapse loss in Alzheimer mouse models”. In: Science 352.6286 (2016), pp. 712–716.

[46] Akash Shah, Uday Kishore, and Abhishek Shastri. “Complement system in Alzheimer’s disease”. In: Int. J. Mol. Sci. 22.24 (2021), p. 13647.

[47] Andrew W Kraft et al. “Attenuating astrocyte activation accelerates plaque pathogenesis in APP/PS1 mice”. In: FASEB J. 27.1 (2013), pp. 187–198.

[48] Ka Chun Tsui et al. “Distribution and inter-regional relationship of amyloid-beta plaque deposition in a 5xFAD mouse model of Alzheimer’s disease”. In: Front. Aging Neurosci. 14 (2022), p. 964336.

[49] Aine M Duffy et al. “Entorhinal cortical defects in Tg2576 mice are present as early as 2-4 months of age”. In: Neurobiol. Aging 36.1 (2015), pp. 134–148.

[50] Maria Bergsland et al. “The establishment of neuronal properties is controlled by Sox4 and Sox11”. In: Genes & Development 20.24 (2006), pp. 3475–3486.

[51] Luming Mu, Lucia Berti, Giacomo Masserdotti, et al. “SoxC transcription factors are required for neuronal differentiation in adult hippocampal neurogenesis”. In: Journal of Neuroscience 32.9 (2012), pp. 3067–3080.

[52] Xinde Zheng, Leah Boyer, Mingji Jin, et al. “Metabolic reprogramming during neuronal differentiation from aerobic glycolysis to neuronal oxidative phosphorylation”. In: eLife 5 (2016), e13374.

[53] Julie E. Cooke and Cecilia B. Moens. “Boundary formation in the hindbrain: Eph only it were simple”. In: Trends in Neurosciences 25.5 (2002), pp. 260–267.

[54] Hantao Shu et al. “Modeling gene regulatory networks using neural network architectures”. In: Nat. Comput. Sci. 1.7 (2021), pp. 491–501.

[55] Shilu Zhang et al. “Inference of cell type-specific gene regulatory networks on cell lineages from single cell omic datasets”. In: Nat. Commun. 14.1 (2023), p. 3064.

[56] Valentine Svensson, Sarah A Teichmann, and Oliver Stegle. “SpatialDE: identification of spatially variable genes”. In: Nat. Methods 15.5 (2018), pp. 343–346.

[57] Shiquan Sun, Jiaqiang Zhu, and Xiang Zhou. “Statistical analysis of spatial expression patterns for spatially resolved transcriptomic studies”. In: Nat. Methods 17.2 (2020), pp. 193–200.

[58] Xiaojie Qiu et al. “Inferring causal gene regulatory networks from coupled single-cell expression dynamics using Scribe”. In: Cell Syst. 10.3 (2020), 265–274.e11.

