## Additional File 1 for "Joint modeling of multi-timepoint spatial observations for time-resolved spatial-unit-specific gene regulatory network inference"

### Supplementary Materials

Yibing Jiang

May, 2025

#### S1. Construction of simulated spatio-temporal datasets

Given an initial GRN matrix, scMultiSim can randomly pick pairs of genes to add new interactions, enlarge interaction strength, remove existing interactions, or weaken interaction strength to generate cell-specific GRNs cell by cell. To induce spatial organization, we used a custom layered layout: we first generate a cluttered, approximately circular set of candidate coordinates, select a bottom-left "origin" location, and then sort cells by Euclidean distance to this origin to form concentric layers. These layers are partitioned into $k$ contiguous spatial domains, and cell types are assigned to domains through a near-diagonal domain–type probability structure so that cells sharing the same GRN class are concentrated within the same domain (with only rare spillover), yielding spatial neighborhoods where nearby cells tend to exhibit similar network patterns. Within each stage, scMultiSim produces cell-specific ground-truth GRNs by gradually rewiring regulatory edges as cells are generated, creating local heterogeneity around a stage-level baseline. To reflect disease-progression continuity and temporal smoothness hypotheses, we chain stages sequentially: stage 1 is initialized from a template GRN provided by the package, while each subsequent stage is initialized by taking a stage-level summary of the previous stage's cell-specific GRNs and applying a controlled sparse rewiring step before simulating the next stage. This ensures that adjacent stages differ incrementally—capturing progressive GRN evolution along a disease axis—while preserving spatio-temporal smoothness (nearby in space and adjacent in time implies more similar GRNs). Temporal precedence is satisfied by construction, since stage $(t)$ is generated using only information from stages $(\leq t-1)$, and no future-stage networks are used to initialize or refine earlier stages.

#### S2. Spatial-guided latent batch correction

During preprocessing, we observed an anomalous global drop in gene expression at 6 months in both genotypes, consistent with a slide-level technical artifact. We corrected this shift using a spatially guided latent batch-correction method inspired by the scLVM concept [1]. To assess whether correction reduced technical suppression without removing biological signal, we compared differential expression between 3M and 6M. This comparison included the original data, the corrected data, and an independent external reference dataset [2]. The corrected data yielded proportions of up- and down-regulated genes that were more consistent with the external reference than those from the uncorrected data (see Fig. 2).

Let $\mathbf{Y} = [y_{ig}] \in \mathbb{R}^{n\times G}$ denote the log-transformed normalized expression matrix, where $n$ is the number of spatial spots, $G$ is the number of genes, and $y_{ig}$ represents the expression of gene $g$ at spot $i$. Define $\mathbf{X} \in \mathbb{R}^{n\times p}$ as the design matrix of known biological covariates, where $\mathbf{X} = [\mathbf{1}, \mathbf{X}_1, \ldots, \mathbf{X}_{p-1}]$, including an intercept column ($\mathbf{1}$) and $p-1$ covariate columns.

We introduce a *guide gene set* $\mathcal{S} \subseteq \{1, \ldots, G\}$ whose biological expression is assumed to be stable across conditions confounded with the slide-level artifact. Their residual variation after covariate regression is assumed to reflect technical artifacts, such as slide-level batch effects, rather than biological signals.

**Residualization of guide genes** For each guide gene $g \in \mathcal{S}$, we model its expression $\mathbf{y}_g = (y_{1g}, \ldots, y_{ng})^\top \in \mathbb{R}^n$ (the $g$-th column of $\mathbf{Y}$) as a linear function of the covariates:

$$\mathbf{y}_g = \mathbf{X}\boldsymbol{\beta}_g + \boldsymbol{\varepsilon}_g, \quad \boldsymbol{\varepsilon}_g \sim \mathcal{N}(\mathbf{0}, \sigma_g^2 \mathbf{I}_n),$$

where $\boldsymbol{\beta}_g \in \mathbb{R}^p$ is the vector of coefficients, $\boldsymbol{\varepsilon}_g \in \mathbb{R}^n$ is the error term with variance $\sigma_g^2$, and $\mathbf{I}_n$ is the $n \times n$ identity matrix. We estimate $\boldsymbol{\beta}_g$ using ordinary least squares (OLS):

$$\hat{\boldsymbol{\beta}}_g = (\mathbf{X}^\top\mathbf{X})^{-1}\mathbf{X}^\top\mathbf{y}_g,$$

assuming $\mathbf{X}$ is full rank. The residual vector for gene $g$ is then:

$$\mathbf{r}_g = \mathbf{y}_g - \mathbf{X}\hat{\boldsymbol{\beta}}_g.$$

Stacking residuals for all guide genes, we obtain the residual matrix $\mathbf{R} = [\mathbf{r}_g]_{g\in\mathcal{S}} \in \mathbb{R}^{n\times|\mathcal{S}|}$, which captures the variation not explained by the covariates, primarily reflecting the technical batch effect.

**Extraction of the latent batch factor** To isolate the dominant technical artifact, we perform principal component analysis (PCA) on $\mathbf{R}$. First, compute the empirical covariance matrix:

$$\mathbf{C} = \frac{1}{n}\mathbf{R}^\top\mathbf{R} \in \mathbb{R}^{|\mathcal{S}|\times|\mathcal{S}|}.$$

Let $\mathbf{C} = \mathbf{V}\boldsymbol{\Lambda}\mathbf{V}^\top$ be its eigendecomposition, where $\boldsymbol{\Lambda} = \mathrm{diag}(\lambda_1, \ldots, \lambda_{|\mathcal{S}|})$ with $\lambda_1 \geq \cdots \geq \lambda_{|\mathcal{S}|} \geq 0$, and $\mathbf{V} = [\mathbf{v}_1, \ldots, \mathbf{v}_{|\mathcal{S}|}]$ contains the corresponding eigenvectors. We extract the first principal component direction $\mathbf{v}_1$, associated with the largest eigenvalue $\lambda_1$, which captures the dominant shared technical variation among guide genes. The latent batch factor is then:

$$\mathbf{u} = \mathbf{R}\mathbf{v}_1 \in \mathbb{R}^n.$$

To ensure numerical stability and interpretability, we standardize $\mathbf{u}$:

$$\mathbf{u} \leftarrow \frac{\mathbf{u} - \mathrm{mean}(\mathbf{u})}{\mathrm{std}(\mathbf{u})},$$

where $\mathrm{mean}(\mathbf{u})$ and $\mathrm{std}(\mathbf{u})$ are the empirical mean and standard deviation of $\mathbf{u}$, respectively. This standardized $\mathbf{u}$ represents the latent batch factor, orthogonal to the covariate space, capturing the slide-level technical artifact.

**Augmented regression and correction** We form the augmented design matrix $\mathbf{Z} = [\mathbf{X} \mid \mathbf{u}] \in \mathbb{R}^{n\times(p+1)}$, incorporating the latent factor $\mathbf{u}$. For each gene $g = 1, \ldots, G$, we model its expression $\mathbf{y}_g$ as:

$$\mathbf{y}_g = \mathbf{X}\boldsymbol{\beta}_g + \mathbf{u}\lambda_g + \boldsymbol{\varepsilon}_g, \quad \boldsymbol{\varepsilon}_g \sim \mathcal{N}(\mathbf{0}, \sigma_g^2\mathbf{I}_n),$$

where $\lambda_g \in \mathbb{R}$ is the gene-specific loading on the latent factor. We estimate the coefficients $(\boldsymbol{\beta}_g, \lambda_g)$ using OLS:

$$\begin{pmatrix} \hat{\boldsymbol{\beta}}_g \\ \hat{\lambda}_g \end{pmatrix} = (\mathbf{Z}^\top\mathbf{Z})^{-1}\mathbf{Z}^\top\mathbf{y}_g,$$

assuming $\mathbf{Z}$ is full rank. The batch-corrected expression for gene $g$ is then:

$$\tilde{\mathbf{y}}_g = \mathbf{y}_g - \hat{\lambda}_g\mathbf{u},$$

and the corrected expression matrix is assembled as $\tilde{\mathbf{Y}} = [\tilde{\mathbf{y}}_1, \ldots, \tilde{\mathbf{y}}_G] \in \mathbb{R}^{n\times G}$.

#### S3. Supplementary Figures

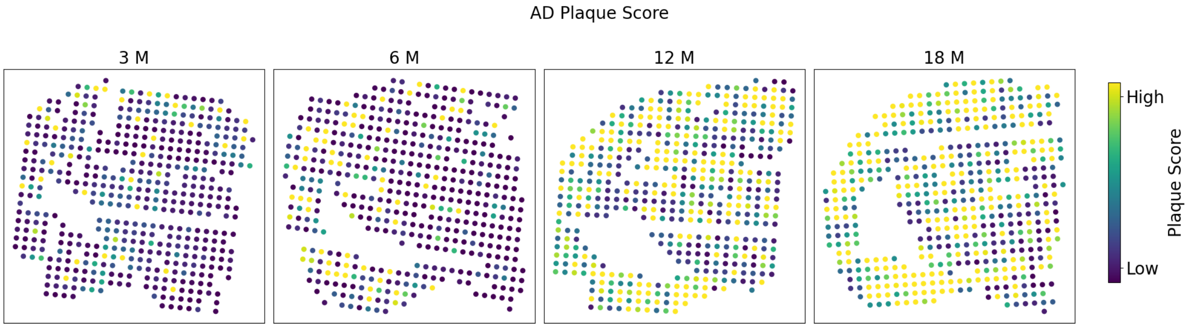


Figure 1: Amyloid plaque burden by spot over time

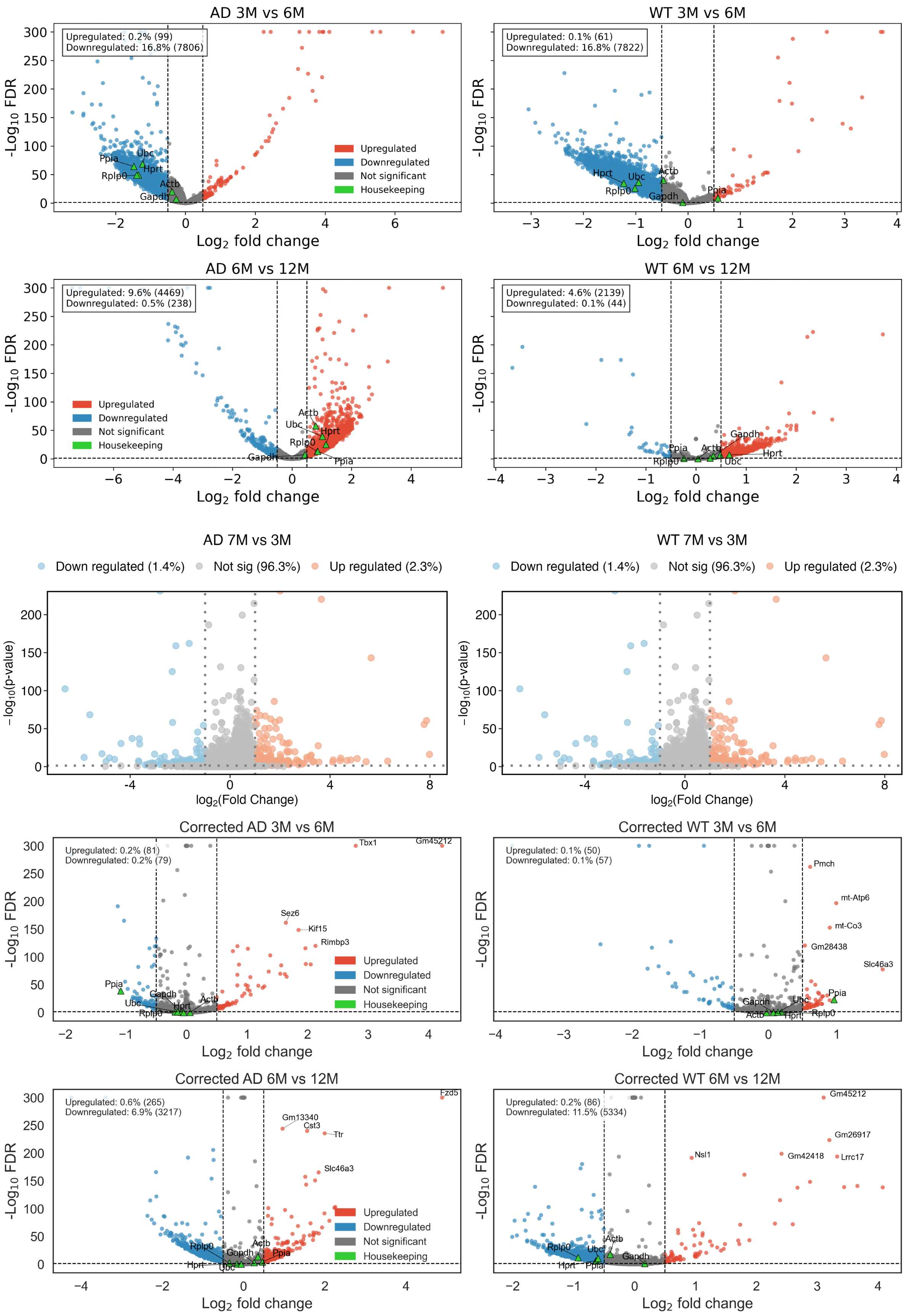


Figure 2: Differential gene expression at 6M in raw data, external dataset, and corrected data

#### References

[1] Florian Buettner et al. "Computational analysis of cell-to-cell heterogeneity in single-cell RNA-sequencing data reveals hidden subpopulations of cells". In: *Nat. Biotechnol.* 33.2 (2015), pp. 155–160. DOI: 10.1038/nbt.3102.

[2] Hongyoon Choi et al. "Spatiotemporal characterization of glial cell activation in an Alzheimer's disease model by spatially resolved transcriptomics". In: *Exp. Mol. Med.* 55.12 (2023), pp. 2564–2575. DOI: 10.1038/s12276-023-01123-9.
